# Assessment of temporal complexity in functional MRI between rest and task conditions

**DOI:** 10.1101/2021.11.20.469367

**Authors:** Amir Omidvarnia, Raphaël Liégeois, Enrico Amico, Maria Giulia Preti, Andrew Zalesky, Dimitri Van De Ville

**Affiliations:** Institute of Bioengineering, Center for Neuroprosthetics, Center for Biomedical Imaging, EPFL, Lausanne, Switzerland; Department of Radiology and Medical Informatics, University of Geneva, Geneva, Switzerland; CIBM Center for Biomedical Imaging, Switzerland; Melbourne Neuropsychiatry Centre, Department of Psychiatry, The University of Melbourne, Melbourne, Australia; Department of Biomedical Engineering, The University of Melbourne, Melbourne, Australia

**Keywords:** fMRI, temporal complexity, multiscale entropy, Hurst exponent, task specificity, graph signal processing

## Abstract

Dynamic models of cortical activity, as measured by functional magnetic resonance imaging (fMRI), have recently brought out important insights into the organization of brain function. In terms of temporal complexity, these hemodynamic signals have been shown to exhibit critical behaviour at the edge between order and disorder. In this study, we aimed to revisit the properties and spatial distribution of temporal complexity in resting state and task fMRI of 100 unrelated subjects from the Human Connectome Project (HCP). First, we compared two common choices of complexity measures (i.e., Hurst exponent versus multiscale entropy) and reported high similarity between them. Second, we investigated the influence of experimental paradigms and found high task-specific complexity. We considered four mental tasks in the HCP database for the analysis: Emotion, Working memory, Social, and Language. Third, we tailored a recently-proposed statistical framework that incorporates the structural connectome, to assess the spatial distribution of complexity measures. These results highlight brain regions including parts of the default mode network and cingulate cortex with significantly stronger complex behaviour than the rest of the brain, irrespective of task. In sum, temporal complexity measures of fMRI are reliable markers of the cognitive status.

## 1 Introduction

In a previous paper [1], we demonstrated reproducible signatures of temporal complexity in resting state networks (RSNs) within a large cohort of healthy subjects. We showed this characteristic of brain function can be reliably quantified through multiscale entropy analysis of resting state fMRI (rsfMRI), and it is distinguishable from head motion. In particular, we reproduced a non-random correlation between temporal complexity of RSNs and higher-order cognition. In this study, we consider a more comprehensive approach by extending our analysis beyond the resting state, considering the complex properties of fMRI, and taking brain structure into account. It is important to be clear about what we mean by the key concept of *complexity* in this paper. Complexity is a balance between order and disorder in different domains. An example of complexity in space (i.e., *spatial complexity*) is a painting which lies between a line with pure regularity and a bunch of colourful curves with pure randomness. An example of complexity in time (i.e., *temporal complexity*) is a song which plays somewhere between a single tone as a pure regular signal and white noise as a pure random signal over time. Of particular interest, temporal complexity has been identified in many dynamical phenomena in nature including the human brain and has been viewed from different perspectives [2]. A common realization of temporal complexity in time series is the existence of *scale-free* dynamics leading to the distribution of similar events across multiple time scales and self-similarity. It speaks to similar states of a signal in the phase space and is associated with self-repetitions of patterns after zooming into the signal again and again. *Multiscale entropy* [3] and *Hurst exponent* [4] are two measures of temporal complexity used in this paper. Multiscale entropy extracts information from coarse grained copies of a signals and provides a representation of signal dynamics at multiple time scales. Hurst exponent quantifies the probability of returning to the signal mean over time and represents the memory of time series.

### 1.1 Temporal complexity and brain dynamics

The concept of complexity has been studied in many real-world phenomena including mechanical systems [5, 6], volcanic eruption [7], climate change [8], earthquakes [9], financial markets [10], biological signals [11, 12, 13, 14, 15, 16], and hemodynamics of the human brain [1, 17, 18, 19, 20]. Of particular interest, brain activity represents a balanced dynamic between local and global functional brain networks at multiple timescales. This *spatiotemporal* complexity originates from a large number of interacting components in the cerebral cortex that are often divided into several subunits themselves with distinctive functional properties. The collective activity of these modules leads to a complex dynamics with a non-linear, non-centralized and self-organized behaviour [21]. Therefore, brain function is a multifaceted phenomenon with different realizations in the time domain. The most important aspects of this complex process include cortical electrical activity measured by electroencephalography (EEG), resulting magnetic fields of the electrical activity of the cortex measured by magnetoencephalogram (MEG), and hemodynamic changes of the brain measured by fMRI. While nonlinear and complex features of EEG and MEG have been vastly studied and confirmed in the literature (e.g., [22, 23]), temporal complexity of fMRI and its link with human behaviour have been less investigated. However, there has been an increasing trend of scientific interest in the temporal complexity analysis of hemodynamic changes in the brain, measured by fMRI, over the recent years [24, 25].

FMRI captures changes of blood flow in the brain through measuring the *blood oxygenation level dependent* (BOLD) signal, which is the basis of RSNs [26]. There is increasing evidence, suggesting that both task-based and rsfMRI have pronounced nonlinear properties which make them navigate through irregular cycles at different frequencies and time scales [27, 28]. It has been hypothesized that (*i*) fMRI is temporally complex, (*ii*) this complexity is brain region dependent and RSN specific, and (*iii*) it is affected by the brain’s dynamical state at rest and during task performance [29]. Self-similarity, one of the realizations of temporal complexity in time series, has also been observed in fMRI in the form of log-linear power spectral density. Ciuciu *et al* [30] showed that fMRI signals have scale-free and multifractal properties during rest and task performance. An important consequence of this finding is that the contribution of different spectral components of fMRI to its dynamic is relatively equal. In other words, missing frequencies of fMRI may be theoretically interpolated from the existing ones. Scale-free dynamics of fMRI is likely affected by mental states. In fact, mental tasks may reduce the self-similarity properties of fMRI [31]. McDonough and Nashiro [17] hypothesized that RSNs may present characteristic complexity patterns. Region-specific properties of fMRI were also studied in [32] where higher complexity was reported in sub-cortical regions such as the caudate, the olfactory gyrus, the amygdala, and the hippocampus, whilst primary sensorimotor and visual areas were associated with lower complexity. Nezafati *et al* [29] confirmed this finding and also, showed that networks exhibit distinct complex properties which may change between resting state and during task performance. Omidvarnia *et al* [1] reproduced the findings of RSN-specific temporal complexity in rsfMRI and lower complexity of sub-cortical regions in contrast to cortical networks across 1000 healthy subjects. They also reported that rsfMRI complexity likely correlates with fluid intelligence. This finding was in line with the hypothesis in [20] where a positive relationship between intelligence and temporal complexity of fMRI was reported. The prefrontal cortex and inferior temporal lobes were amongst brain regions with the strongest relationship between high fMRI complexity and high intelligence. These studies, to name but a few, suggest that evaluation and analysis of fMRI fluctuations can provide insight into functional brain networks, dynamics of brain structure and human behaviour. There is evidence that the temporal complexity of brain function supports different aspects of human behaviour and cognition [33]. Perturbed complexity across cortical areas may contribute to a range of brain diseases including epilepsy [34], Alzheimer’s disease [35], and schizophrenia [36]. Time-varying changes of functional brain networks are likely related to the fine balance between efficient information-processing and metabolic costs in the brain [37]. Characterization of the neural correlates of temporal complexity in brain function can shed light on how interactions between cortical regions are temporally organized and has the capacity of leading to imaging-based biomarkers of brain function in health and disease. A necessary step towards this goal is to assess the reliability and reproducibility of fMRI complexity aspects in healthy subjects.

Different measures have been used for temporal complexity analysis of fMRI including time-resolved graph theory measures [34], entropy measures [1, 17, 20, 29, 34], and self-similarity measures [30, 31, 38, 39]. Among these, Hurst exponent [4] and multiscale entropy [3] have found a wide acceptance due to their straightforward interpretation and fairly simple implementation. A crucial step towards an appropriate understanding of temporal complexity measures of fMRI is to ensure that these quantifications are able to differentiate between background fluctuations and structured changes in the fMRI time series. This step is done through statistical hypothesis testing where a given null hypothesis expressing randomness will be invalidated [40]. A suitable null distribution must preserve all properties of the data except the one which is under investigation. Surrogate data analysis is a widely used non-parametric approach for developing such null distributions. This approach is based on the shuffling of a single feature or a set of features in the data, while the other fundamental features are kept intact. In a complicated dynamical system with a combined complexity in time and space such as the human brain, appropriate null distributions must take both the dynamics and the underlying structure into account. Phase shuffling is an effective way of generating surrogates of fMRI when the cross-correlations between brain regions need to be preserved [41]. However, it fails to consider the anatomical basis of fMRI into account. Establishing a brain structure-informed statistical inference for temporal complexity analysis of fMRI is still an open question. A solution to this challenge is through graph signal processing [42] where fMRI time series of different brain regions are projected onto the underlying structural brain graph and randomization is done in the brain graph space instead of the time domain [43]. It can help to better understand the structural substrates of temporal complexity in brain hemodynamics and its spatial distribution across brain areas.

In this study, we aim to perform an independent assessment on the most commonly reported aspects of temporal complexity in fMRI during rest and task engagement. We use two measures of temporal complexity, i.e., Hurst exponent and multiscsle entropy, and compare them in the context of fMRI analysis. First, we validate the monofractal feature of brain hemodynamics during task and rest using Hurst exponent of fMRI in a population of healthy subjects from the Human Connectome Project (HCP) [44]. Second, we assess task specificity of fMRI complexity during task engagement and resting state. To this end, we perform a pair-wise support vector machine (SVM) analysis on the Hurst exponent of fMRI. Third, we examine the hypothesis of complexity suppression of brain hemodynamics due to task engagement. Fourth, we investigate the agreement between spatial distribution of fMRI complexity, obtained by Hurst exponent and multiscale entropy. Finally, we perform statistical testing on fMRI complexity through a brain structure-informed surrogate technique which generates null distributions by randomizing the human connectome. We look into the mathematical properties of this technique in detail and its consequences for our analysis. Figure 1 illustrates the procedure of performing temporal complexity analysis and graph surrogate generation on a typical fMRI dataset, as adapted in this study.

**Figure 1:**
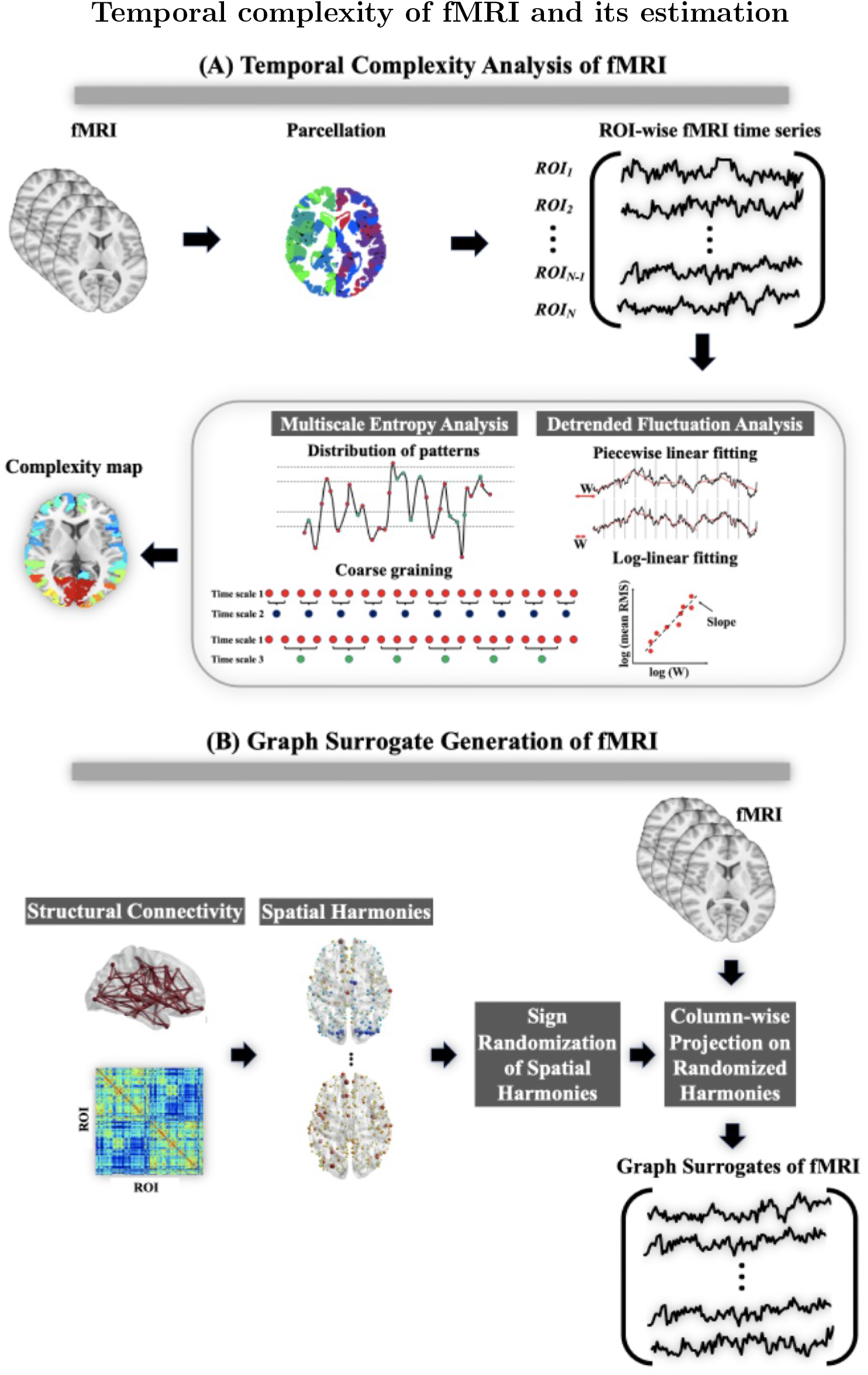
An overview of the paper. (A) Complexity is a balance between order and disorder in different domains. An example of spatial complexity is a painting which lies between pure regularity (such as a line) and pure randomness (such as a bunch of colourful random curves). An example of the temporal complexity is a song which is somewhere between a single tone as a pure regular signal and white noise as a pure random signal over time. (B) Two common realizations of temporal complexity in time series include the distribution of similar events across multiple time scales and self-similarity. The former speaks to the similar parts of a signal in the phase space and the latter is associated with self-repetitions after zooming into the signal again and again. (C) Multiscale entropy and Hurst exponent are two measures of temporal complexity used in this paper. Multiscale entropy extracts sample entropy from coarse grained copies of a signals and provides a representation of signal dynamics at multiple time scales. Hurst exponent, measured here through detrended fluctuation analysis, quantifies the probability of coming back to the signal mean over time. It is done by checking the log-linear distribution of slopes of piece-wise linear trends in the time series. (D) In this paper, we investigate three aspects of temporal complexity in fMRI datasets of 100 unrelated subjects. We look into the spatial distribution of fMRI temporal complexity and sensitivity of different brain regions to this balanced dynamic. We then check task specificity of fMRI temporal complexity at rest and during task engagement. We finally look into the link between fMRI temporal complexity and brain structure by adapting a graph surrogate testing method based on the combination of brain function and structure.

## 2 Materials and methods

### 2.1 Data and preprocessing

In this study, we the rest and task fMRI and diffusion-weighted scans of 100 unrelated subjects (ages 22-35) from the Human Connectome Project (HCP) 1200-release [44]. Each subject underwent a number of fMRI recording sessions including four rsfMRI runs, seven task-based fMRI runs, and a diffusion MRI run. Each fMRI dataset had a voxel size of 2×2×2 millimetres and the repetition time (*T_R_*) of 720 milliseconds in a 3-T Siemens Skyra scanner. We utilised two rest runs with left-right phase encoding as well as four task runs with a minimum length of 3 min or 250 *T_R_*’s, i.e., Language, Motor, Social, and Working Memory. The other three tasks, i.e., Gambling, Emotion, and Relational, were shorter than 3 minutes and therefore, excluded from the analysis. Note that each task-based fMRI recording had a specific task design with different number of conditions and trials (Table 1). See [48] for the description of the fMRI tasks. We included the rsfMRI datasets with left-right phase encoding only due to the known issue of asymmetric drop-out between left-right and right-left phase encoding of rest runs in the HCP database [45] and its potential impact on ask specificity analysis in section 2.3. Since the length of rsfMRI in all subjects was considerably longer than all task fMRI recordings (14.4 min versus 3-4 min), we used the first 399 *T_R_*’s of rest runs in order to make the extracted complexity measures comparable. The fMRI datasets were preprocessed using SPM8 through a procedure described in [47]. A parcellation mask [46] was used to parcellate the grey matter into 360 cortical regions of interest (N_*ROI*_=360). The corresponding diffusion-weighted datasets were preprocessed through the steps outlined in [47] and used to extract the structural connectivity matrix of each subject.

**Table 1:** List of fMRI runs for each subject in the HCP

| Run | Session | $N_{TR}$ | Length in *minutes* | No. of conditions | No. of trials |
| --- | --- | --- | --- | --- | --- |
| 1 | Rest 1 LR | 399 | 4.8 | - | - |
| 2 | Rest 2 LR | 399 | 4.8 | - | - |
| 5 | Language | 305 | 3.67 | 2 | 11 |
| 6 | Motor | 273 | 3.29 | 5 | 10 |
| 7 | Social | 263 | 3.17 | 2 | 5 |
| 8 | Working Memory | 395 | 4.74 | 8 | 8 |

### 2.2 Temporal complexity analysis of fMRI

One of the most common ways to measure scale invariance in time series is by using the Hurst exponent *H* [4]. It provides insight about self-similarity of signals, i.e., their tendency to revert back or cluster to a long term equilibrium. It also determines whether there is a predominant time-scale or frequency component in the underlying dynamical process. The value of *H* varies between 0 and 1 where *H* ≤ 0.5 implies short-memory or fast return to the mean (such as white noise), and *H* ≥ 0.5 represents long-memory or a trending behaviour with random turning points. The value of *H* = 0.5 represents a random walk whose time points have no correlation with their past values. Scale-free signals have a long-memory, because all of their time scales and spectral components contribute equally to their dynamics. It leads to a power law relationship in the spectral power of scale-free signals in the form of *P*(*f*) ∝ *f^β^* where *P*(*f*) is the power spectral density at frequency *f*, and *β* is a non-zero positive real number referred to the *spectral exponent*. For some scale-free processes such the fractional Brownian motion, there is a theoretical relationship between the spectral exponent *β* and the Hurst exponent *H* via the equation *β* = 2*H* – 1. In this study, we used detrended fluctuation analysis (DFA) [49] to estimate the Hurst exponent and log-linear line fitting to the power spectral density function to estimate the spectral exponent of fMRI time series. The DFA algorithm has been used in the previous studies for the analysis of fMRI fractality [31, 50]. Given the direct link between the Hurst exponent and signal entropy [51], we also looked into the complex behaviour of fMRI using multiscale entropy [3]. This measure is based on the sample entropy [52] at several time scales of a signal **x** (here, an fMRI time series at a particular ROI). See Appendix A for a detailed description of multiscale entropy.

### 2.3 Task specificity of fMRI complexity

In order to test the dependency of complex properties of brain hemodynamics to mental states, we used a set of linear Support vector machines (SVMs) in order to evaluate the separability of the Hurst exponent over 4 rest runs and 4 task runs of fMRI. SVM analysis is a supervised learning method which is widely used for performing classification and regression studies using fMRI datasets. A binary SVM takes the data points of two classes and estimates a hyperplane in the feature space that best separates the class labels. We extracted ROI-wise complexity measures of *N_subj_*=100 subjects and 6 fMRI runs. For each pair-wise comparison between the two fMRI runs, we considered each subject as an observation and the brain maps as feature vectors of size *N_ROI_* × 1 where *N_ROI_*=360. It yielded 100 feature vectors of size *N_ROI_* × 1 for each task. Here, we investigated if mental tasks can be discriminated in a population of healthy subjects using ROI-wise Hurst exponent of fMRI. This pair-wise SVM analysis led to two symmetric accuracy matrices of size 6 × 6 for the sensitivity and specificity of pair-wise classifications. The classification accuracy of each SVM classifier was calculated via leave-one-out cross validation and its hyper-parameters were optimized through grid search.

### 2.4 Spatial distribution of fMRI complexity across grey matter

To be able to interpret the complexity measures of fMRI across brain regions, one must perform statistical testing. In this study, we generated *N_Surr_*=100 surrogate fMRI datasets for each subject and each fMRI run through a *graph signal processing* framework which combines the brain structure and function in order to generate surrogates of fMRI whose null distribution preserves the anatomically-defined linear representation of the data [42, 43]. Our motivation for adapting this technique was to incorporate the underlying anatomical aspects of fMRI in the significance testing step, an important information which is usually neglected in the fMRI complexity analysis studies. As discussed in Appendix B, the graph surrogate method: (*i*) preserves linear properties of each fMRI channel, (*ii*) preserves temporal relationship between fMRI time points, i.e., temporal correlation, (*iii*) randomizes functional connectivity, (*iv*) randomizes the spatial variation of scale-free dynamics of fMRI at each single ROI, (*v*) randomizes the spatial variation of the scale-free dynamics between ROI pairs.

In order to obtain group-level maps of brain complexity, we applied binomial testing on the subject-specific brain maps. First, each individual map of complexity was thresholded at a significant level of *α_subj_*=0.01 in order to obtain a binary map. Then, the binomial distribution *P*(*n*) of having *n* detections was used at each ROI to examine the significant number of suprathreshold regions across subjects at the significant level of *α_group_*=0.001. It was equivalent with 10 *successes* (i.e., number of suprathreshold regions) for a population of 100 subjects,. The results were corrected for multiple comparisons for the number of regions and fMRI runs tested (360 × 6).

## 3 Results

### 3.1 FMRI represents complex behaviour during rest and task

As seen in Figure 2, normalized group mean power spectral density of all fMRI recording sessions (4 fMRI runs and 4 task runs) show log-linearity in the frequency domain. This feature was more pronounced in rsfMRI datasets and the spectral exponents were RSN-specific with the default mode network (DMN), frontoparietal network (FP), and dorsal attention network (DA) presenting the highest exponents in most cases (Figure 2-B). The *β* exponents of different RSNs, however, were shown to be task-dependent. For example, Social task led to the highest spectral exponent across subjects at the DA network or rest runs led to the highest exponents at the DMN and FP network. A striking observation was related to the existence of dominant peaks in the log-linear power spectral density functions of task fMRI in contrast to rsfMRI (Figure 2-A). In order to rule out the influence of the task designs in this spectral feature of task fMRI recordings (Figure 2-C), we regressed out the task timings from the data and checked spectral log-linearity of the residuals only. As Figure 2-D illustrates, the peaks still remain in the log-linear power spectral density functions of task fMRI residuals, although they are slightly suppressed. It suggests that these spectral peaks are independent from the task design and likely explain why the Hurst exponent is generally smaller during task engagement and larger at rest [30, 31, 40].

**Figure 2:**
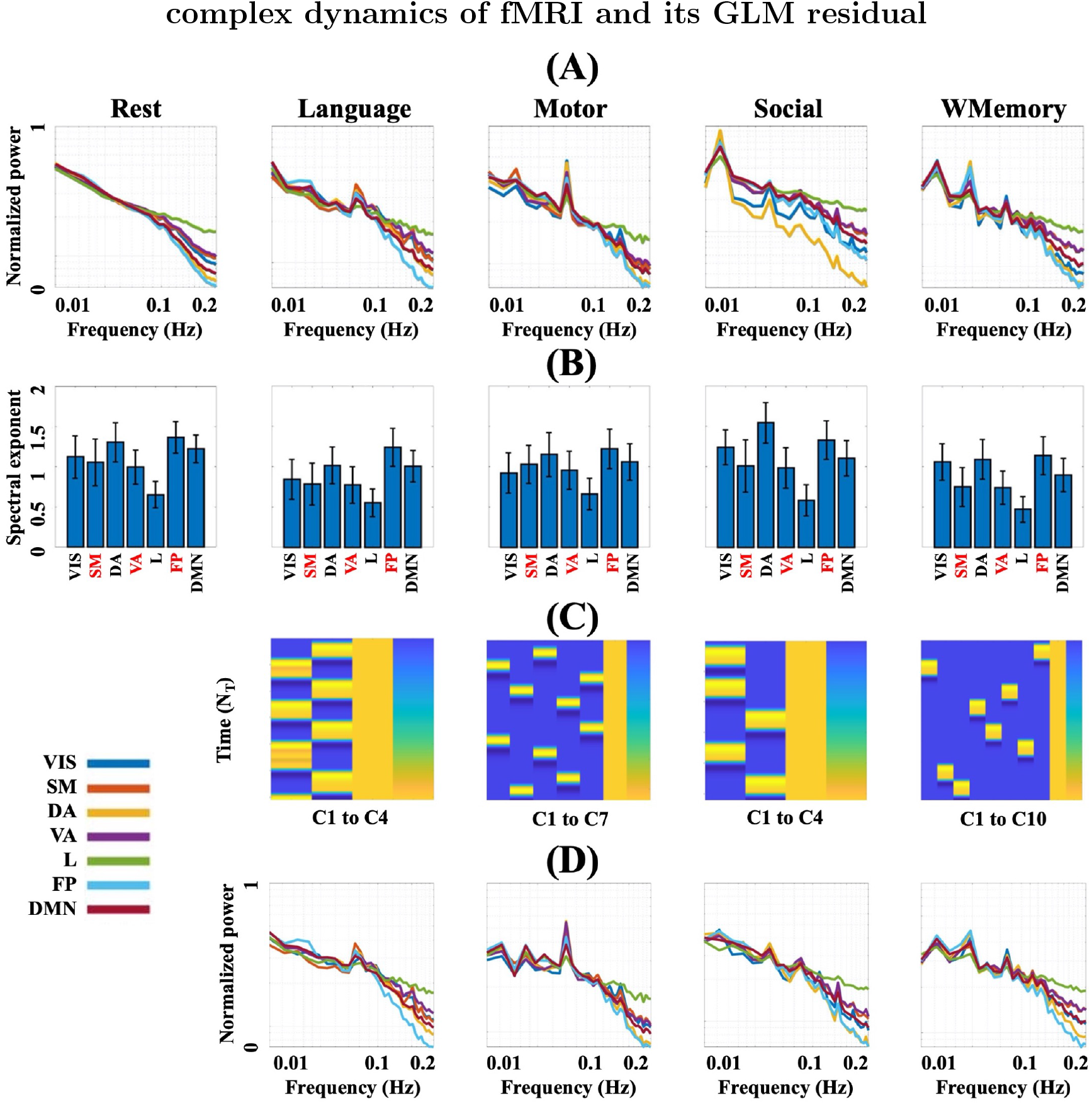
Normalized power spectra of RSNs averaged over subjects and the corresponding *β* exponents, estimated within the frequency band of 0.02-0.2 Hz. Abbreviations: VIS=Visual, SM=Somatomotor, DA=Dorsal attention, VA=Ventral attention, L=Limbic, FP=Frontoparietal, DMN=Default mode network, numbered C=Condition.

### 3.2 Task engagement suppresses complex dynamics of fMRI

As summarized in the specificity and sensitivity matrices of Figure 3, mental tasks can be classified with high accuracy (up to 100%) using the Hurst exponents of their associated fMRI recordings. Also, ordering of the slice acquisition seems to have a significant impact on the Hurst exponent of rsfMRI (note the high classification rates between left-right and right-left rsfMRI recordings in Figure 3-C). Here, each dataset has been characterized as a set of *N_subj_* feature vectors of size *N_ROI_* × 1 where *N_subj_* = 100 and *N_ROI_* = 360. The distribution of Hurst exponent across brain areas in different rest and task fMRI runs (brain maps of Figure 3-A) suggests that the spatial profile of complex dynamics in fMRI heavily depends on the task engagement and resting conditions. As evident in the histograms of individualized mean Husrt exponents across brain areas in Figure 3-B, each task results in a regionally specific reduction in dynamic complexity, an observation in line with previous findings in the literature [31]. It establishes a framework for comparing the *task burden* in subjects. For example, the Hurst exponent histograms suggest that the working memory task is likely the most *demanding* brain state in the population of this study, because its associated histogram covers the lowest interval of self-similarity.

**Figure 3:**
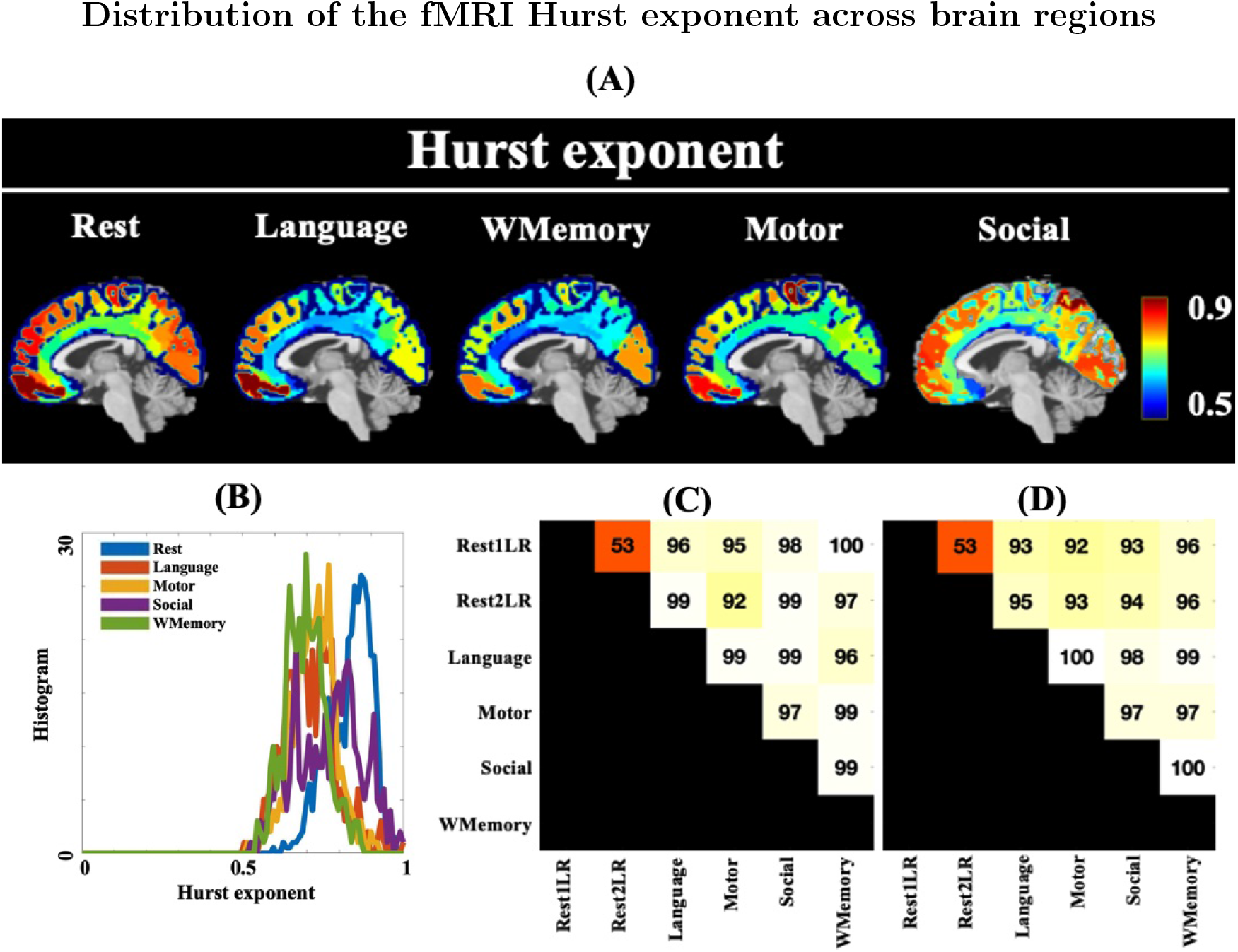
(A) Spatial distributions and histogram of the Hurst exponent across brain regions (averaged over subjects). The brain maps of 4 rest runs have been averaged. (B) Histograms of the group mean Hurst exponent over 360 brain regions for 4 task runs and 4 rest runs (averaged). Classification accuracy of pair-wise comparison of mental tasks using binary SVM classifiers with linear kernel: (C) sensitivity of the Hurst exponent (87.1%±13.2%), and (D) specificity of the Hurst exponent (85.3%±14.1%). Abbreviations: WMemory=Working memory, Rest1LR=First Rest run with left to right slicing, Rest2LR=Second Rest run with left to right slicing.

### 3.3 Complex dynamics exist in the brain structural-functional coupling

Figure 5 demonstrates the logarithmic spectral power and multiscale entropy patterns of the projected fMRI datasets onto brain structure at rest and during task engagement. The lowest spatial harmony in brain connectome (zero frequency in Figure 2-A) is associated with the mean of the fMRI temporal correlation matrix. As the figure shows, the distribution of spatial energy across brain connectome follows a log-linear relationship. However, despite what we observed for the spectral power of fMRI in Figure 2-A, a considerable difference between the power spectral distributions of graph signals extracted from different tasks and rest sessions is not evident. This can also be observed in the multiscale entropy patterns and complexity indices of brain graph signals in Figure 2-B, C which are quite comparable over four tasks and the average rest. In particular, there is an inflection point in the scatter plot of complexity indices (Figure 2-C) associated with the color transition in the multiscale entropy patterns of Figure 2-B from dark blue (low randomness) to green (high randomness). These *elbow* or *knee* points are another indicative of log-linearity in the complexity domain of brain structure-function.

**Figure 4:**
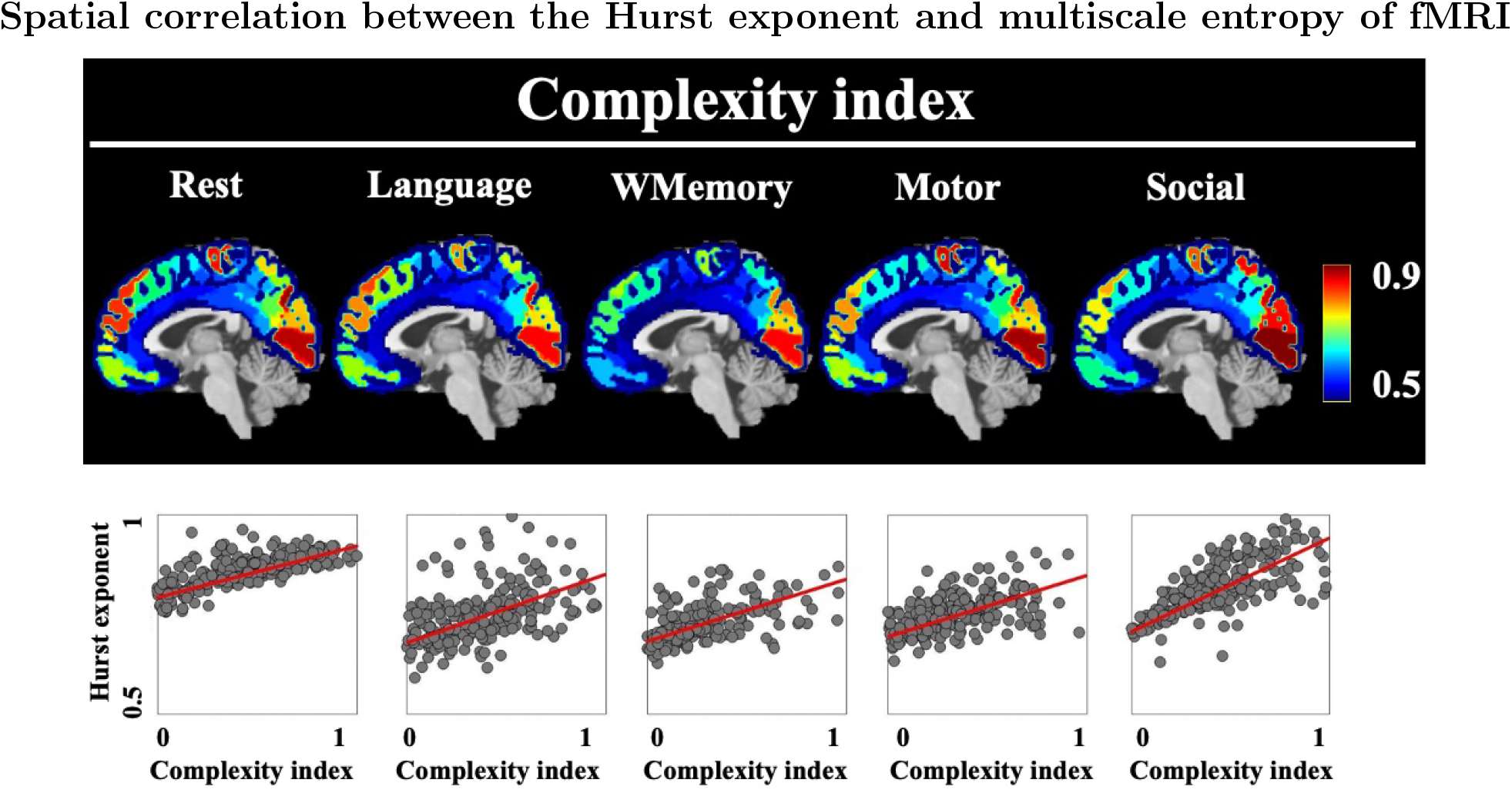
Upper row: Spatial distributions of the entropy-based complexity index across brain regions (averaged over subjects). Lower row: Joint distribution of the Hurst exponent and complexity index extracted from rest and task fMRI datasets, averaged across all subjects. The brain maps of 4 rest runs have been averaged. Abbreviation: WMemory=Working memory.

**Figure 5:**
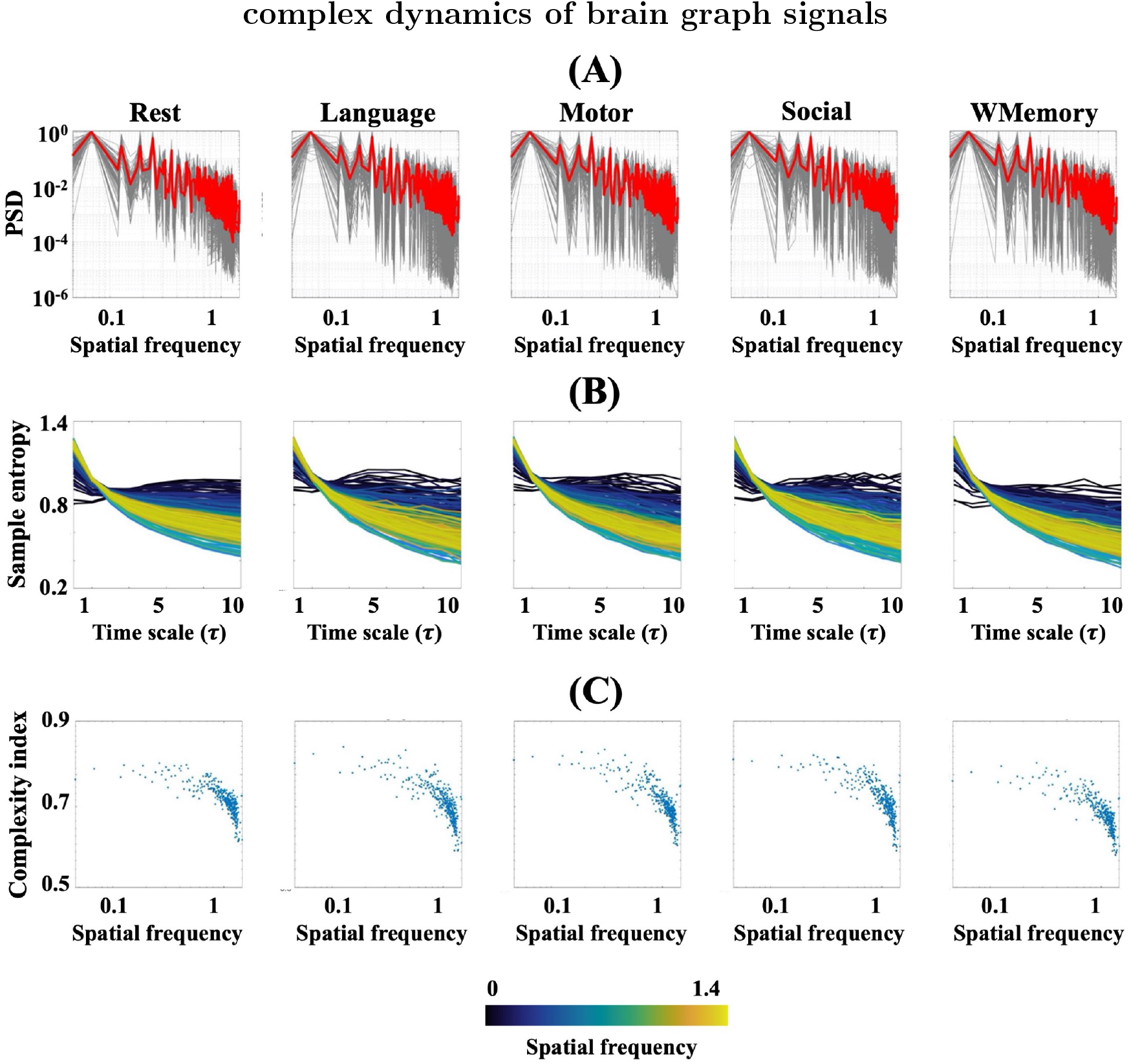
(A) Logarithmic plots of the power spectral density functions of brain graph signals (i.e., projection of the fMRI data at rest and task onto brain structure) versus brain spatial harmonies. The plots of 4 rest runs have been averaged. Each gray curve belongs to a certain subject and the red curves represent group mean. All curves have been normalized to 1. (B) Multiscale entropy patterns of the graph signals, colour-coded by their associated brain spatial harmony. (C) The complexity indices associated with the multiscale entropy curves of (B).

### 3.4 Spatial patterns of complex dynamics in fMRI

Figure 6-A illustrates the group-mean spatial patterns of complexity index across brain areas for the average rest runs and 4 tasks. Also, Figure 6-B provides a binary equivalent of Figure 6-A where each suprathreshold brain region is presented by a bar and RSNs have been highlighted with different transparent colors for illustration purpose. All maps were thresholded at the subject-level *p*-value of 0.01 and family-wise error corrected at the group-level *p*-value of 0.01 using the graph surrogate data generation method introduced in [43]. Also, Table 2 summarizes the contribution of 7 RSNs in the suprathreshold brain regions of Figure 6. In all fMRI runs, frontoparietal network involved the highest percentage of brain regions with significant complex dynamics (up to 70.8% of engagement). The next mostly engaged RSNs in all runs were the dorsal attention network with a maximum cover of 63% in the Social task followed by the default mode network with a maximum cover of 47.8% in the average Rest run. Several regions across dorsal lateral prefrontal cortex (*DLPFC*) showed up as the most frequently observed areas with the highest complexity across all tasks and rest runs. These regions included *8Av, 9m, 8BL, 10d, 9a*, and *p10p*. Other most repeated regions included: left *RSC* (anterior cingulate cortex), left *TE1a* (middle temporal gyrus), *PGi* (temporo-parieto occipital junction), *PGs* (inferior parietal cortex), and right *45* (inferior frontal cortex). The limbic network represented the least complex regions for all task and rest runs. See [46] for the anatomical description of these brain labels.

**Figure 6:**
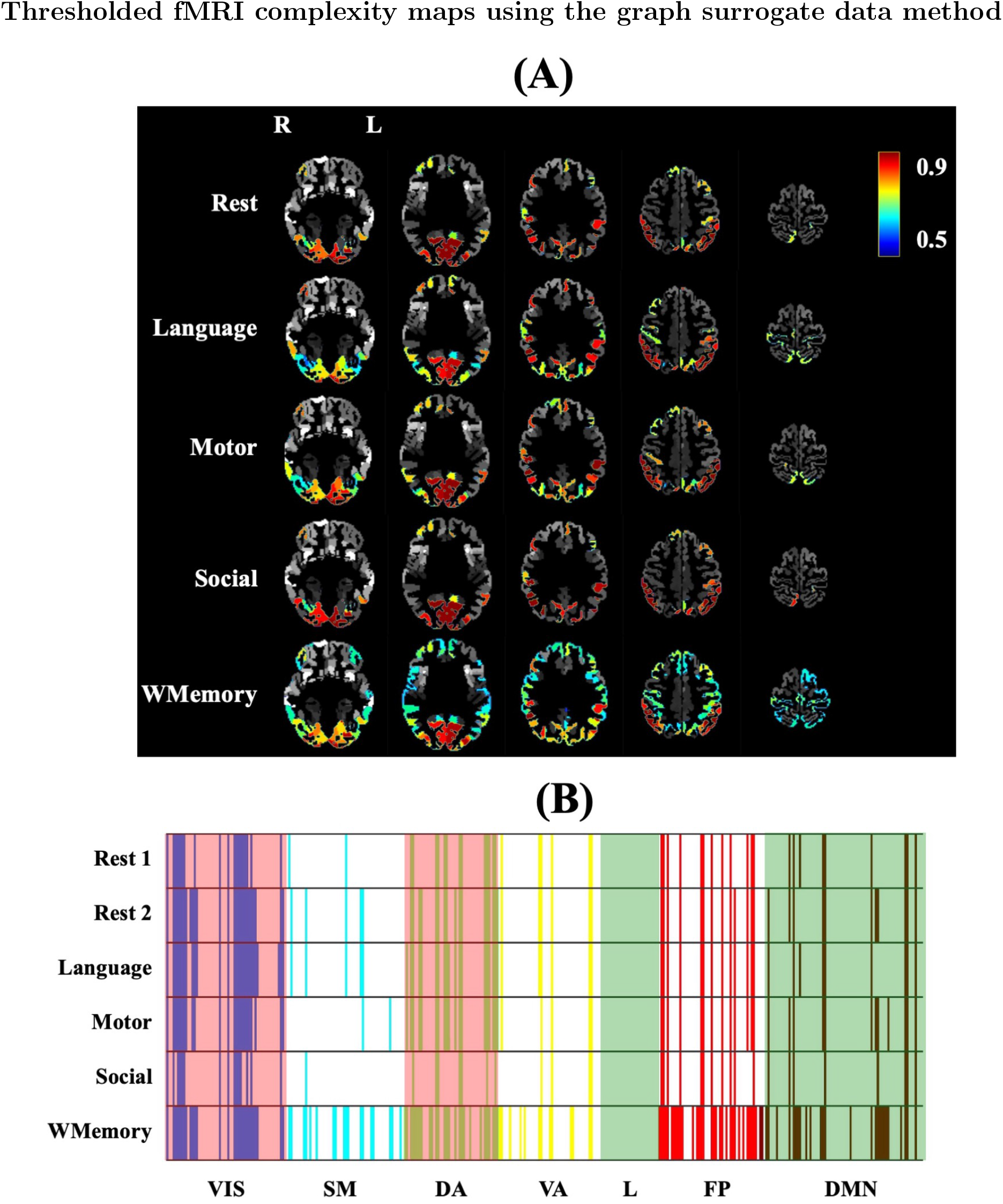
(A) Spatial distribution of group-mean fMRI temporal complexity across brain areas for 4 task runs and the average rest run. All maps have been thresholded using the graph surrogate data generation [43] at the subject level *p*-value of 0.01 and family-wise error corrected at the *p*-value of 0.01. (B) The equivalent binary representation of the brain maps in panel (A) where each suprathreshold brain region has been illustrated as a bar and the RSNs have been highlighted with different transparent colors for an easier visual perception. Abbreviations: VIS=Visual, SM=Somatomotor, DA=Dorsal attention, VA=Ventral attention, L=Limbic, FP=Frontoparietal, DMN=Default mode network, WMemory=Working memory. Note that the label ’L’ in panel (A) represents ’Left’ and not ’Limbic’.

**Table 2:** Percentage of suprathreshold ROIs in 7 RSNs after graph surrogate testing of brain complexity maps in Figure 6. Abbreviations: VIS=Visual, SM=Somatomotor, DA=Dorsal attention, VA=Ventral attention, L=Limbic, FP=Frontoparietal, DMN=Default mode network.

| Task name | VIS | SM | DA | VA | L | FP | DMN |
| --- | --- | --- | --- | --- | --- | --- | --- |
| Rest 1 LR | 32.8% | 3.7% | 28.3% | 12.5% | 0% | 25.0% | 11.5% |
| Rest 2 LR | 46.6% | 9.3% | 36.9% | 12.5% | 0% | 27.1% | 14.1% |
| Language | 48.3% | 9.3% | 41.3% | 12.5% | 0% | 27.1% | 11.5% |
| Motor | 41.3% | 3.7% | 41.3% | 10.4% | 0% | 27.1% | 15.4% |
| Social | 25.9% | 1.9% | 15.2% | 8.3% | 0% | 22.9% | 10.2% |
| Working Memory | 48.3% | 33.3% | 58.7% | 27.1% | 0% | 66.7% | 35.9% |

## 4 Discussion

This study reinforces the existence of complex dynamics in brain function [29] and provides further evidence for the hypothesis of distinct complexity features in human behaviour and cognition [31]. Our results suggest that: (*i*) task-based and rsfMRI signals exhibit temporal complexity, inferred by Hurst exponent and multiscale entropy, (*ii*) rest and task periods of brain function can be distinguished from each other with high accuracy based on their temporal complexity profiles, (*iii*) cognitive load can suppress the complex dynamics of fMRI in contrast to the resting state, (*iv*) spatial distribution of Hurst exponent and entropy-based complexity index in fMRI are highly correlated, and (*v*) the frontoparietal network and default mode network represent maximal complex behaviour compared to the rest of the brain regardless of the mental state.

Fluctuations of neural activity in the brain are observed on the scale of milliseconds in single-cell spiking to the order of seconds in BOLD changes [53]. In contrast to the traditional neuroscientific view which would mostly consider this variability as random disturbances and measurement noise, new investigations have discovered many systematic patterns of information in neural variability over multiple temporal and spatial scales [3, 54, 55]. Neural variability at the BOLD level not only reflects inter-subject differences such as behavioural traits, but it also captures within-subject changes such as dynamical states of the brain and cognitive load [53]. An important feature of neural variability in the brain is its temporally complex behaviour. Although temporal variability and temporal complexity are closely related, these two concepts are not necessarily the same. In other words, temporal complexity always exhibits variability, but a variable process is not necessarily complex. A complex pattern of brain activity is rich in information over time and represents a balanced dynamic between order and disorder within brain networks or between different brain regions [1, 17]. Recent theories in neuroscience have proposed that temporal complexity of brain function is likely associated with information processing in the brain [17, 55]. The first theory [56] indicates that a healthy brain retains an optimal degree of instability which allows it to enter to *different dynamical states* and sample the external world. In this way, the brain learns how to optimize responses to environmental stimuli. The second set of theories [57] argue that a moderate level of randomness is necessary in the neural system because it increases the probability of *neuronal firing* in subthreshold neurons. In contrast to the second proposal, a third view [58] is that temporal complexity of brain function can enhance or suppress the likelihood of *neural synchrony* between cortical areas. All of these theories speak to the direct relationship between temporal complexity of brain function and information transfer across neural populations, cortical regions and functional networks.

Considering the key role of complex processes in brain mechanisms, one would expect to find realizations of temporal complexity in brain hemodynamics as well. To investigate this possibility, it is crucial to quantify temporal complexity of neural variability. There are several interpretations of signal complexity across the literature with tendency towards specific aspects of dynamical systems or specific signal domains. The most observed aspects in neural variability and brain function include: (*i*) determinism, (*ii*) randomness, and (*iii*) self-similarity/fractality/complex dynamic. Determinism, in its nonlinear sense, is mostly studied using chaos theory which is *the qualitative study of unstable aperiodic behaviour in deterministic nonlinear dynamical systems* [59]. It models a signal through a set of differential equations with time-constant parameters. Sometimes determinism is not in the signal dynamics *per se*, but it is in its statistical properties. Randomness speaks to the arbitrary changes of these properties such as mean and standard deviation in time. A process is self-similar, fractal or complex if it repeats the same statistical characteristics across many temporal scales. Intuitively, the more a signal is self-similar, the more its long-term memory increases and the less it generates new information by passing time. Based on this speculative view on temporal complexity, the measures of neural variability can be categorized into three classes [53]: variance-based, frequency-based, and entropy-based. Although all of these three classes have been used in previous studies, the latter has found wide acceptance for the temporal complexity analysis of fMRI. This is mainly because entropy measures do not rely on distributional assumptions (unlike variance-based measures), and are not restricted by sinusoidal signal waveforms (unlike frequency-based measures). In this study, we utilized two measures for temporal complexity analysis of fMRI in order to cross-check the fMRI complexity analysis results: multiscale entropy as an entropy-based measure which evaluates similar patterns of information throughout signals and the Hurst exponent as a variance-based measure which quantifies the memory of signals and their tendency to return to their mean values. Multiscale entropy has been shown to be sensitive to RSN-specific hemodynamics, and reproducible across healthy subjects [1, 17]. The direct relationship between multiscale entropy and self-similarity [51] makes it a good candidate for investigating the complex properties of fMRI and comparable with the Hurst exponent of fMRI.

It is important to note that the existence of self-similarity and fractality in BOLD signals has been subject to discussion over the past decades, mainly due to the limited temporal resolution of fMRI and sluggishness of the hemodynamic responses in the brain [60]. A fundamental question here is that even assuming the presence of temporal complexity in brain hemodynamics, whether the recorded BOLD signal inside a MRI scanner is able to adequately capture it. To address this question, the following issues should be considered: (*i*) low temporal resolution of the measured BOLD changes with a typical *T_R_* value in the scale of hundred milliseconds to seconds, (*ii*) the use of short-length fMRI time series (less than 200 *T_R_*’s) in some studies, (*iii*) potentially manipulative impact of preprocessing steps and residual scanning artifacts on the nonlinear dynamics of fMRI, (*v*) upon agreement on the complex dynamics of fMRI, whether it is monofractal or multifractal [30, 61]. It has been shown that non-fractal time series with inadequate memory can still exhibit log-linear spectral power and be falsely identified as complex processes [60]. Therefore, one may argue that the observed complex behaviour of fMRI is not biological, but it simply originates from the *signal aspects* of fMRI such as its inadequate length in the previous studies. In fact, it has been shown that the accuracy of fMRI fractal analysis can be affected by the number of brain volumes [61]. Despite these technical difficulties, several studies have supported the hypothesis of a power law distribution for the fMRI power spectrum over the frequency band of 0.01 Hz to 0.1 Hz [30, 50, 62, 63]. There is also evidence that the scale invariance temporal dynamics in fMRI is of a multifractal nature. However, monofractal analysis of fMRI would be a safer option, because it quantifies the complex structure of fMRI at lower magnifications in contrast to multifrctal analysis and therefore, needs lower signal length and temporal resolution [30, 61]. For this reason, we chose to perform the monofractal analysis of fMRI in this study using Hurst exponent *H*.

As Figure 4 shows, a linear association across regions was evident between the Hurst exponent and the area under multiscale entropy curves. Although the two measures operate at different ranges and utilize different methodologies to quantify temporal complexity, their spatial agreement across brain regions is very high (Pearson correlation above 0.8). In particular, both measures represent maximal values across the visual cortex, anterior cingulate cortex and parts of DMN. The distinction between the dynamics of rsfMRI and task fMRI is more apparent in the Hurst exponent brain maps (Figure 3) in contrast to the entropy-based complexity index brain maps (Figure 4). However, we performed statistical testing on the multiscale entropy brain maps only, because the Hurst exponent of real-world signals (such as fMRI) can be affected by the choice of the estimation method making it difficult to reproduce the results by other researchers. On the other hand, estimation of multiscale entropy and the associated complexity index is straightforward and easy to replicate.

The magnitude of *H* across brain areas in our study (Figure 3) is comparable with the previous findings showing a typical range of ≈0.5-1 for *H* [31, 64, 65]. The spatial extent of fMRI temporal complexity in our results is maximal across the frontoparietal and default mode networks and minimal across deep brain areas such as the limbic network (see Figure 3 and Figure 4), in line with the previous studies of fMRI [1, 31]. The considerable overlap between the analysis results of this study with the existing literature suggests that the fMRI signal length (minimum of 263 *T_R_*’s - see Table 1) has been enough to replicate the previous hypotheses about temporal complexity of brain function. In fact, a previous study has showed that valid Hurst exponents can be obtained from fMRI time series as brief as 40 sec (≈ 56 *T_R_*’s) through the DFA algorithm [31]. Also, it implies that the fMRI preprocessing steps have not significantly manipulated the true dynamics of fMRI datasets in this study. According to our results, the spatial patterns of Hurst exponent in fMRI are highly correlated with the associated patterns of entropy-based complexity index. It indicates that the two measures of fMRI temporal complexity converge to a similar outcome even though their computation is completely different. The sampling rate of fMRI (*T_R_*) has to be much higher than its highest frequency component for temporal complexity analysis, so that it can capture the true dynamics of the hemodynamic process [61]. This requirement is met for the HCP fMRI datasets with a *T_R_* of 0.72 sec, because its sampling frequency, i.e., 1.34 Hz, is much higher than the upper-band of fMRI frequency content, i.e., 0.1 Hz.

As Figure 3-A and Figure 4-A illustrate, brain regions with the highest temporal complexity are commonly located across the frontoparietal, visual, anterior cingulate cortex and default mode networks in all task and rest fMRI runs. On the other hand, the sub-cortical areas and mid-cingulate cortex are found to have the least complex dynamics. The brain maps illustrated in the figures also suggest temporal complexity of some regions such as the motor areas is task-dependant, meaning that their complex property seems to vary over different tasks. High temporal complexity and Hurst exponent of a given brain region or RSN mean that the associated fMRI signals are temporally redundant and predictable. Therefore, highly complex regions/networks are likely responsible for the process of internal stimuli with low surprise and high adaptability. However, external stimuli such as sensory inputs and auditory inputs may reduce adaptability of brain dynamics, increase surprise, and suppress temporal complexity of brain function. An exception would be the visual network which shows high temporal complexity and high average Hurst exponent, likely due to the larger capacity of visual areas in contrast to the other networks and brain regions. It is important to note that despite the presence of complex dynamic in fMRI, the relationship between fMRI temporal complexity and intrinsic functional connectivity is scale-dependant. It has been shown that the weighted sum of functional links to a given brain node, referred to as *functional connectivity strength* or FCS, is associated with the functional significance of that node in support of information transfer across the brain [66]. The link between FCS and temporal complexity of rsfMRI has been shown to be scale-dependant [1, 17]. In particular, an inverse relationship has been reported between FCS and temporal complexity of RSNs at fine time scales of multiscale entropy (*τ* ≤ 5 at a *T_R_* of 0.72 sec equivalent with time periods shorter than 3.5 sec to 4 sec), while it turns to a proportional relationship at coarse time scales (*τ* > 6 or time periods greater than ≈ 4 sec). This scale-dependant relationship varies for different RSNs. For example, frontoparietal and default mode networks represent the highest correlation between resting state FCS and temporal complexity at fine scales and lowest correlation at coarse scales, while it is the opposite for somatomotor, sub-cortical and visual networks [1, 17]. On the other hand, fine time scales of fMRI have been associated with the dynamics of local neural populations, whilst the coarse time scales are likely related to the long-range functional connections [17]. Altogether, these observations would suggest that in order to obtain a comprehensive picture about complex dynamics of fMRI, one must consider the anatomical locations (i.e., spatial distribution) of brain regions. It speaks to the necessity of appropriate spatiotemporal methods for significance testing of fMRI temporal complexity which can account for brain structure and function at the same time.

Surrogate data testing is a powerful method for characterizing the statistical properties of time series. In this approach, we want to compare the measure of interest extracted from the original data, i.e., the alternative hypothesis *H*_1_, to the distribution of the same measure obtained from a large number of surrogate data, i.e., the null hypothesis *H*_0_. In the context of fMRI temporal complexity, the null hypothesis could be that fMRI time series at different brain regions are generated by some non-complex processes and their functional relationships are also non-complex. If the complexity indices of fMRI signals fall within the null distribution *H*_0_, it means that these indices only rely on the statistical properties preserved under the null *H*_0_. Otherwise, we can reject the null hypothesis and interpret fMRI temporal complexity as revealing statistical properties beyond *H*_0_. A critical question here is how to specify the null hypothesis of scale-free dynamics in fMRI and how to remove this signal feature of interest from the original data in order to generate surrogates [67, 68]. In this study, we chose brain graph randomization technique [43] which shuffles the power spectral density of fMRI, while keeping its underlying anatomical properties. As analytically discussed in section B.3 and Appendix B, the surrogate data in this study preserves temporal correlation of fMRI. However, functional connectivity, spatial variation of complex dynamics, and spatial variation of the complex dynamics of functional connectivity are randomized in the surrogate time series. Therefore, complex dynamics of brain ROIs with suprathreshold complexity indices is significantly different from chance level and likely have a key role in moderating the information transfer across the whole brain. According to Table 2 as well as section 3.4, the supporting role of brain regions is maximal within the frontoparietal, visual, and default mode networks such as *DLPFC* and *PGi*, regardless of the resting state or task engagement. This finding is in line with the previous studies showing the dominant role of these brain areas in internal processing and facilitation of information exchange in functional brain networks.

A number of caveats need to be considered in interpreting the results of this study. From the technical perspective, one should be aware of the limitation of multiscale entropy in capturing multivariate relations between fMRI time series at multiple ROIs. The standard version of multiscale entropy used in this study treats ROI-wise fMRI signals as a set of individual and independent time series. However, mean fMRI time series at different ROIs are often statically dependent and correlated due to the smearing effect of hemodyanamic changes in the brain and possible effect of fMRI preprocessing steps such as spatial smoothing. Therefore, it is plausible to use the multivariate versions of temporal complexity measures for fMRI data analysis [69, 70]. A systematic comparison between univariate and multivariate measures of complex dynamics and temporal complexity remains for our future work. Another consideration should be given to the nature of BOLD signals as an indirect measure of neural activity and its reduced amount of information it carries, in contrast to other direct measurements with higher temporal resolution such as local field potentials. Although fMRI has a greater capacity to cover larger brain areas than localized measurements of neural activity and can provide large scale information about complex properties of brain dynamics, its temporal complexity has to be treated as an indirect property of brain function. Having said that, the dependency of fMRI temporal complexity to different mental states (as illustrated in Figure 2 and its association with the hubs of information transfer in the brain is an evidence that the BOLD signals still preserve some significant aspects of neural populations [29]. In this study, we assumed monofractality in fMRI and compared entropy-based complexity index of fMRI with the classical Hurst exponent at different ROIs. It would be informative to check the possible links between multiscale entropy and multifractality of fMRI using longer fMRI datasets and higher temporal resolutions during rest and task engagement.

## 5 Conclusion

Temporal complexity is a reproducible aspect of fMRI during rest and task engagement. This feature of brain function is task specific and can be suppressed by cognitive load. FMRI complexity is a discriminative feature between rest and task in the brain functional domain, but not in the brain structural domain. A brain structure-informed statistical testing of fMRI complexity reveals several areas with suprathreshold temporal complexity within the frontoparietal, visual, and default mode networks.

## Author contributions

## Acknowledgement

AO acknowledges support through the Eurotech Postdoc Programme, co-funded by the European Commission under its framework programme Horizon 2020 (Grant Agreement number 754462). RL acknowledges support by the Swiss National Centre of Competence in Research - Evolving Language (grant number 51NF40 180888). EA acknowledges financial support from the SNSF Ambizione project ”Fingerprinting the brain: network science to extract features of cognition, behaviour and dysfunction” (grant number PZ00P2-185716). MGP was supported by the CIBM Center for Biomedical Imaging, a Swiss research center of excellence founded and supported by Lausanne University Hospital (CHUV), University of Lausanne (UNIL), Ecole Polytechnique Fédérale de Lausanne (EPFL), University of Geneva (UNIGE) and Geneva University Hospitals (HUG). The primary fMRI data in this study was provided by the Human Connectome Project, WUMinn Consortium (1U54MH091657; Principal Investigators: David Van Essen and Kamil Ugurbil) funded by the 16 National Institutes of Health (NIH) institutes and centres that support the NIH Blueprint for Neuroscience Research; and by the McDonnell Center for Systems Neuroscience at Washington University.

## Appendix

### A Multiscale entropy analysis

Multiscale entropy analysis [3] is based on the calculation of sample entropy [52] at multiple time scales. Sample entropy treats each short piece of **x** as a *template* to quantify a conditional probability that two templates of length *m*, which are similar to within a tolerance level *r*, will remain similar when *m* becomes *m* +1. Note that self-matches are not considered in calculating this conditional probability. A template 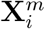 is defined as^1^:

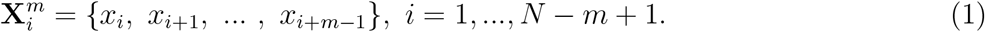

where *N* is the number of time points in **x** and *m* is the embedding dimension parameter. Two templates 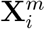 and 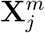 are considered as neighbours if their Chebyshev distance 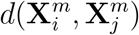 is less than a *tolerance* parameter *r*. It leads to an *r*-neighbourhood conditional probability function 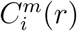 for any vector 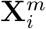 in the *m*-dimensional reconstructed phase space:

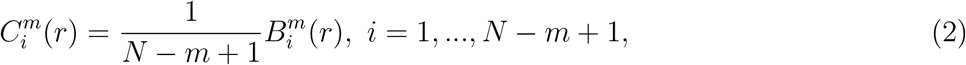

where 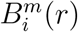 is given by:

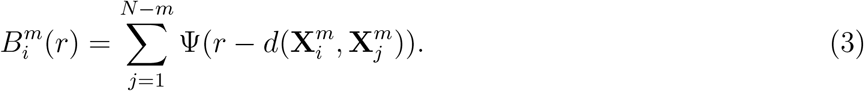

Sample entropy is then given by:

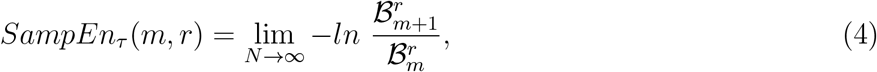

where 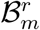 is the average of 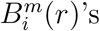 over all templates:

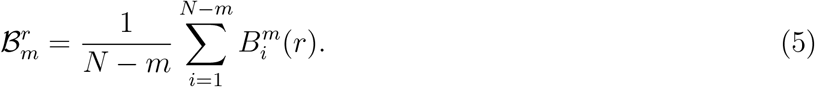

Sample entropy is always non-negative, but it can also become undefined. It is important to multiply the tolerance parameter *r* by the standard deviation of **x** to account for amplitude variations across different signals [52]. Multiscale entropy extracts sample entropy after *coarse-graining* of the input signal **x** at a range of time scales *τ*. A coarse-grained vector **x**(*τ*) = {*x_i_*(*τ*)} is defined as:

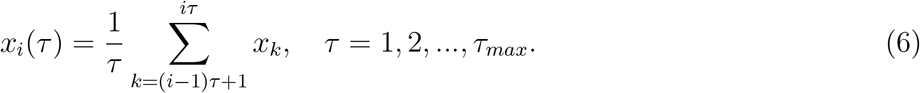

In this study, we used these parameter values: *m* = 2, *r* = 0.5, and *τ_max_* = 10. We reduced the dimensionality of multivariate multiscale entropy patterns to a single value by calculating the area under each multi-scale entropy curve over all scales, divided by the maximum number of scales (i.e., *τ_max_*). It leads to a *complexity index M_i_*, for each parcellated rsfMRI dataset, approximated with the normalized area under the multivariate sample entropy values across multiple time scales as follows [71]:

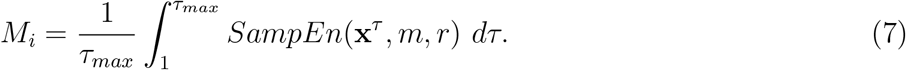

Figure 7 demonstrates an example of two signals with different dynamics and distinct temporal complexity profiles: white noise (in red color) and Brown noise (in blue color). According to the definition of temporal complexity in this paper, white noise is not a complex signal because it represents an extreme case of disorder with no specific pattern of structured information in the time and frequency domains. It is apparent from the uniformly distributed spectral power of white noise (Figure 7-B), a decreasing pattern of multiscale entropy (Figure 7-D), and low complexity indices (Figure 7-E). On the other hand, Brown noise represents a complex dynamic reflected in its log-linear power spectral density function (Figure 7-C) leading to significantly larger complexity indices than white noise (Figure 7-E, associated with the larger areas under the entropy curves in Figure 7-D).

**Figure 7:**
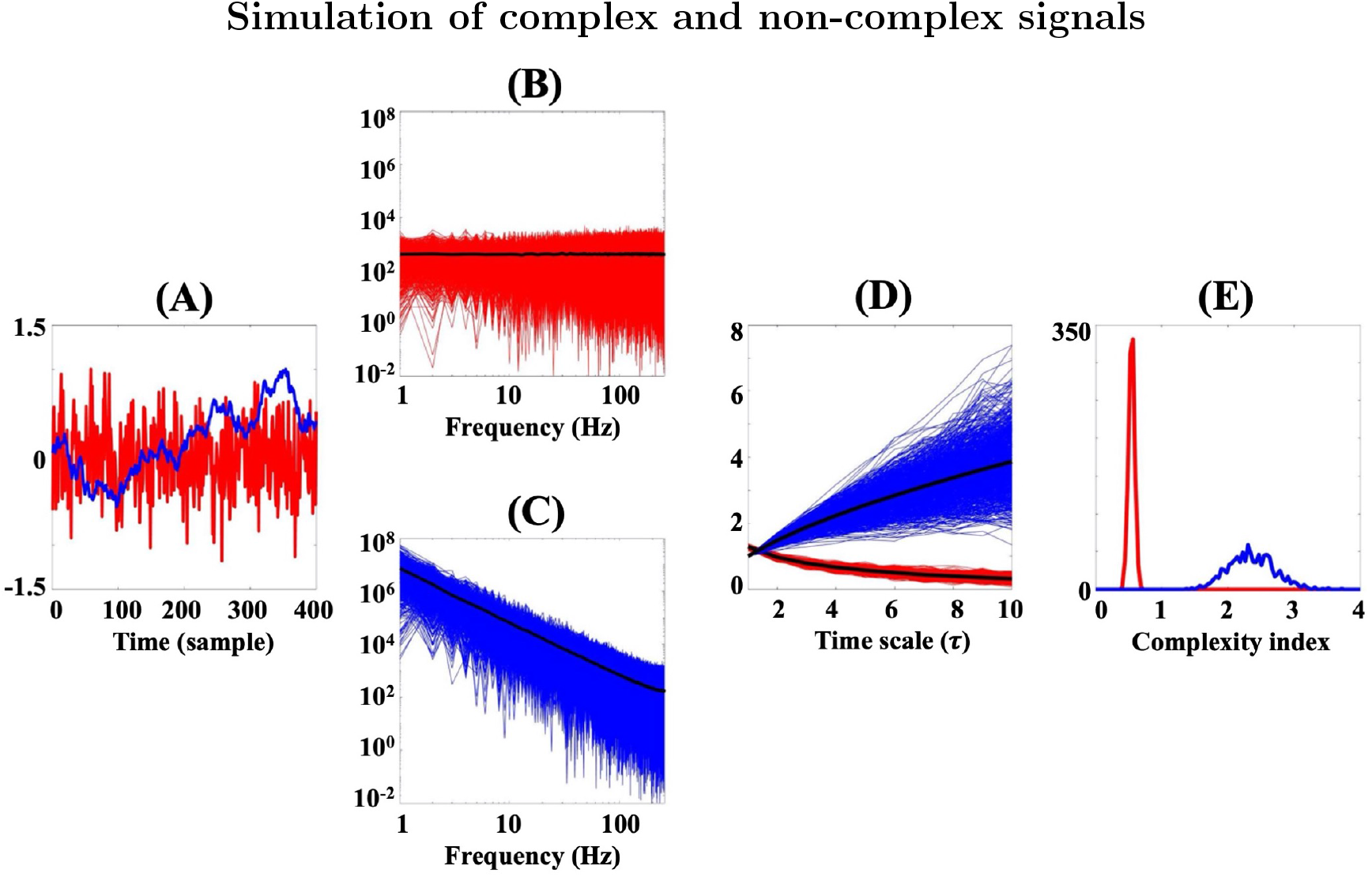
Simulation of white noise and Brown noise

### B Graph surrogate method for fMRI complexity analysis

In this section, we explain a surrogate data method, proposed in [43], which we believe is a relevant technique for generating null distributions in hypothesis testing of fMRI complexity. This approach uses the graph signal processing framework [42] in order to combine brain structure and function. Let 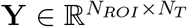 be a parcellated rsfMRI dataset from *N_ROI_* brain regions of interest (ROIs) which have been uniformly sampled at *N_T_* time points. We then assume that 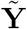 is the graph surrogate form of **Y** and its associated fMRI temporal correlation matrix 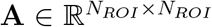, that is, 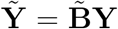 (see also Eq. 13) [43]. Also, let **B** be a matrix whose columns are the spatial eigenvectors of **A**.

#### B.1 Combining brain structure and function

In order to model the spatiotemporal correlates of fMRI, we consider a basis set 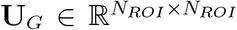 which contains the spatial harmonies of brain function and another basis set 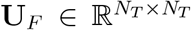 which models the dynamics of fMRI using a Fourier transform with *N_T_* bins. Joint projection of **Y** into the brain structural and spectral domains can be modelled as:

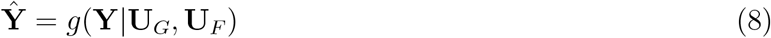

where 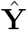 is a *N_ROI_ × N_T_* matrix and *g* is a nonlinear mapping from the time domain into the joint complexity-structure domain. For the sake of simplicity, we assume that *g* is a linear function and also, the structural and spectral domains of brain function are separable. In this case, Eq. (8) is written as:

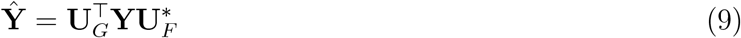

where 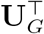 is the transpose of **U**_*G*_ and * denotes the complex conjugate operator. This general formulation considers the spatial and spectral properties of brain function through **U**_*G*_ and **U**_*F*_, respectively. Assuming **U**_*F*_ as an identity matrix, one can obtain the representation of graph signal processing for fMRI datasets.

#### B.2 Spatial harmonies of brain structure

In order to obtain the structural basis set of brain function (i.e., **U**_*G*_ in Eq. 9), one can incorporate a *brain structural graph* from the corresponding diffusion MRI [42]. This graph can be characterized as *G* = (*V, E*) where *V* is a set of *N_ROI_* vertices and *E* is a set of weighted edges associated with a symmetric and real-valued adjacency matrix 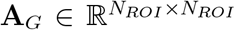. Each element in **A**_*G*_ represents the number of white-matter pathways between two brain regions. The symmetric normalized Laplacian matrix of **A**_*G*_ is then used to form a *structural space* onto which brain function is projected:

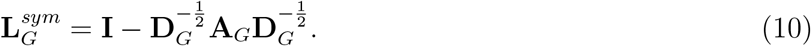

where 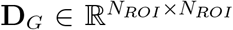 is the identity matrix and 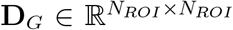 is the degree matrix of **A**_*G*_, a diagonal matrix whose non-zero elements are defined as:

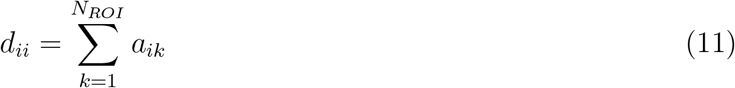

where *a_ik_* is the *ik*^th^ element of **A**_*G*_. According to the spectral graph theory [42, 72], spatial harmonies of the fMRI temporal correlation matrix *G* can be obtained through eigendecomposition of 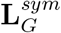 as follows:

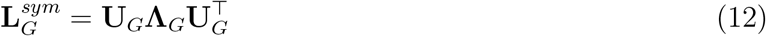

where the diagonal matrix **Λ**_*G*_ includes the eigenvalues (or spatial harmonies) of 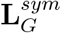 and **U**_*G*_ contains the corresponding spatial eigenmodes. Given that **A**_*G*_ is real-valued and symmetric, both **Λ**_*G*_ and **U**_G_ will be real-valued. The matrix **U**_*G*_ constitutes an *N_ROI_*-dimensional *brain structural space* or *structural space*, in short. The assumption here is that brain structure is fixed in time. Therefore, all elements of **U**_*G*_ are time-invariant. The left-side multiplication of 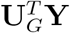 in Eg. 9 is usually referred to as *graph signals* in the literature [42].

#### B.3 Graph surrogate generation

We used the graph surrogate data generation method in [43] to develop null distributions of fMRI complexity. This technique is based on the idea of sign randomization in brain structural eigenmodes. Let 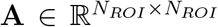 be the fMRI temporal correlation matrix associated with the vectorized fMRI data 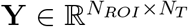. The spatial eigenmodes of **A** could be obtained through graph signal processing as the columns of a square and orthogonal matrix 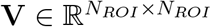 where **VV**^*T*^ = **I** and **I** is the identity matrix.

One can randomize the eigen matrix **V** by keeping the absolute value of its elements, while shuffling of the signs of eigenvectors (i.e., the signs of all elements across a column are either flipped or not). In this case, a surrogate form of **Y** with a similar size, called hereafter 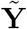, is obtained as [43]:

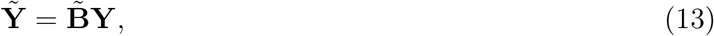

where 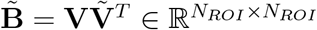. Each element of 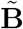 can be written as:

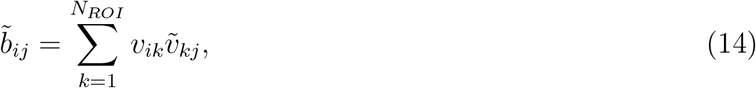

where *v_ik_* is the *ik*^th^ element of **V** and 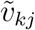 is the *kj^th^* element of 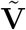. In Eq. 13, 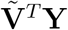 randomizes the fMRI matrix **Y** in the structural brain graph domain and left-side multiplication to **V** takes the randomized graph signals back to the functional domain. Note that in the case of 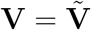 (i.e., if there is no shuffling in the graph domain), we have:

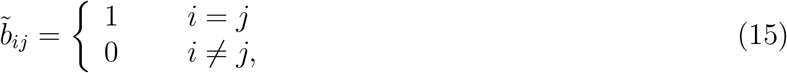

while in the case of graph shuffling, we have:

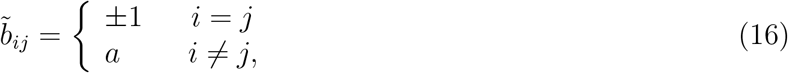

where 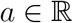 is an unknown real number.

#### B.4 Regarding linearity

The surrogate data matrix 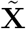 in Eq. 13 can be also written as:

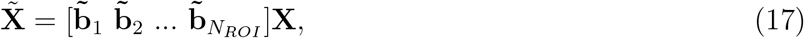

where 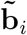 is the *i*^th^ column in 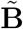. By assuming the index *j* as a temporal argument in Eq. 17, the *i*^th^ element of 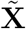 at time *t* is given by:

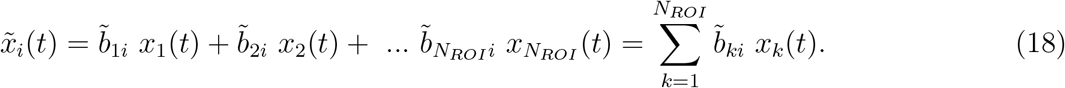

It implies that each element of the surrogate data 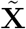 at time *t* is a linear random combination of its corresponding fMRI time point over all ROIs. Therefore, the graph surrogate method [43] preserves the linear relationships between original fMRI time points.

#### B.5 Regarding functional connectivity

Now, let us compute the *spatial* covariance matrix of 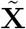, referred to as 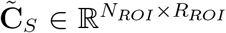, in order to get an insight about inter-regional cross-correlation surrogate time series as follows:

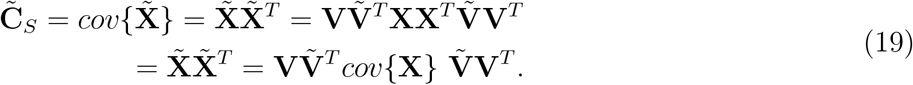

The matrix 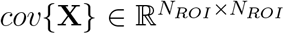 is equivalent with the *functional connectivity* extracted from fMRI data **X**. Since 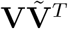 and 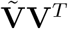 are not necessarily identity matrices, 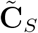 and *cov*{**X**} will not be the same. Therefore, the graph surrogate method in [43] randomizes functional connectivity.

#### B.6 Regarding the fMRI temporal correlation matrix

To unravel the pair-wise relationship between the time points of graph surrogates, a similar crosscorrelation analysis can be done by simply transposing 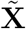 in Eq. 19. It leads to the *super adjacency* matrix 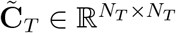 as follows:

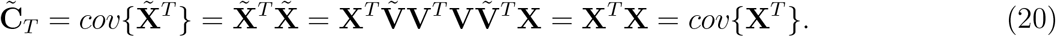

This is based on the fact that both **V** and 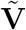 are orthogonal, so the terms 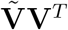 and 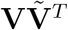 are identity matrices. Since 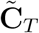 is independent from 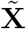, it is not changed throughout the surrogate generation procedure. It suggests that the fMRI temporal correlation matrix embedded in the surrogate data matrix 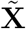 is preserved through the method described in [43].

#### B.7 Regarding the complex properties

The complex properties of a multivariate time series can be studied by looking into the log-linearity of its auto- and cross-spectral power in the frequency domain. A log-linear pattern of spectral power means that its frequency components have been distributed according to an exponential law *S*(*f*) = 1/*f^β^* where *β* is referred to as the *spectral exponent*. This parameter is related to the *memory* of the underlying signal measured by *Hurst exponent H*, varying between 0 and 1. A value near 0 reflects short memory (i.e., quickly coming back to the signal mean or noise-like behaviour), while a value close to 1 represents long memory (i.e., slow return to the signal mean or a smooth dynamic). In the spacial case of fractional Brownian motion, an exponent of *H* = 0.5 leads to a random walk. Also, Hurst exponent and spectral exponent are linked through a simple relationship: *β* = 2*H* + 1.

In order to check the complex properties of graph surrogates in 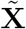, one can use the linear expansion in Eq. 18 and obtain the Fourier transform of 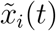 as follows:

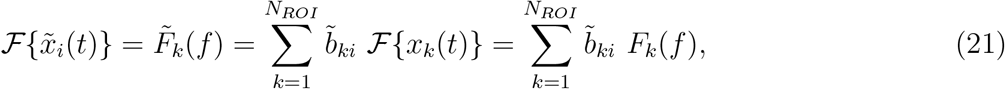

where *x_k_*(*t*) is the original fMRI time series at the *k*^th^ ROI. The power spectral density function of 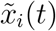 is then obtained as:

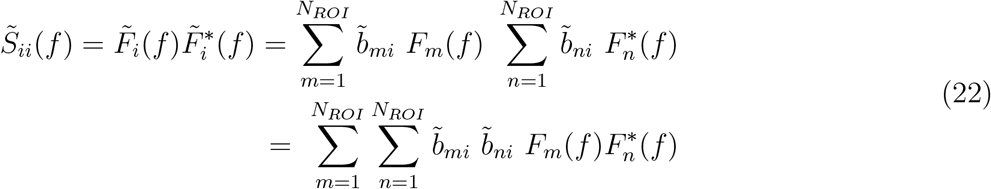

Each single multiplication term 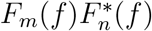 in Eq. 22 is either equal to the power spectral density of individual surrogate time series (i.e., *S_mm_*(*f*)) or coherency between pairs of surrogate signals (i.e., *S_mn_*(*f*)) as follows:

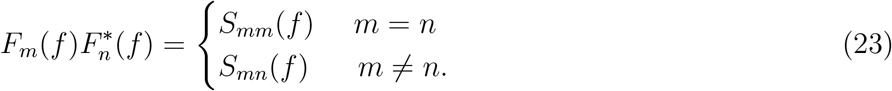

Therefore, the spectral power of each surrogate time series in 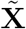 is a random linear combination of all spectral powers and all possible pair-wise coherencies between original fMRI time series across ROIs. Now, let us assume that there is some level of complex dynamics within the original fMRI time series, that is, the spectral power and coherency across a sub-group or all temporal dimensions of **X** follow power law:

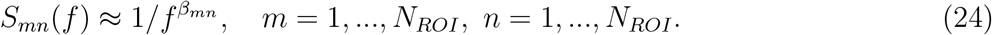

With this assumption, it is clear from Eq. 22 that the power spectral density functions of surrogate data 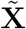 (i.e., 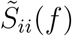, *i* = 1, …, *N_ROI_*) do not necessarily preserve the complex properties of the original fMRI data at individual ROIs or interaction between regions. If there is no variation in complex properties across brain regions (i.e., if the spatial distribution of spectral exponents is the same across all regions), the Hurst exponent of surrogates would be the same as the original data. This is an example where the complex properties of the original dataset are completely preserved by the graph surrogate data method [43]. In practice, however, the fMRI temporal correlation matrix **A** imposes a non-uniform distribution of the complex properties across regions which can be picked up by the graph surrogate method.

Note that the coherency between different graph surrogates (i.e., 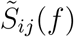, *i, j* = 1, …, *N_ROI_*, *i* ≠ *j*) gives a *shuffled version of functional connectivity* in the time domain. To show this, we take advantage of the *Cross-Correlation* and *Wiener–Khinchin* theorems [ref] outlining the reciprocal relationship between correlation and coherence in the time and frequency domains. The Wiener-Khinchin theorem states that for 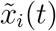, the power spectral density 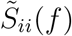 is equal to the Fourier transform of its auto-correlation function 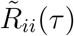:

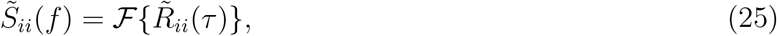

where *τ* denotes the delay and 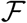 is the Fourier transform operator. Also, Cross-Correlation theorem indicates that the correlation between surrogates 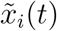 and 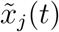 in the time domain is equivalent with their coherence in the frequency domain:

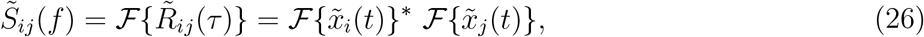

where 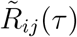 is the cross-correlation of 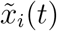 with 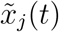 at the delay *τ*, and * denotes the complex conjugate operator. It suggests that the graph surrogate method [43] tends to remove spatial variation of possibly complex properties of the original data at each single fMRI time series, it preserves the temporal relationships, but randomizes their functional connectivity.

## Footnotes

1 In all equations, scalar variables are in normal font, while vector variables are in bold.

